# Distinct Microbiomes Underlie Divergent Responses of Methane Emissions from Diverse Wetland soils to Oxygen Shifts

**DOI:** 10.1101/2024.12.19.629461

**Authors:** Linta Reji, Jianshu Duan, Satish Myneni, Xinning Zhang

## Abstract

Hydrological shifts in wetlands, a globally important methane (CH_4_) source, are critical constraints on CH_4_ emissions and carbon-climate feedbacks. A limited understanding of how hydrologically driven oxygen (O_2_) variability affects microbial CH_4_ cycling in diverse wetlands makes wetland CH_4_ emissions uncertain. Transient O_2_ exposure significantly stimulated anoxic CH_4_ production in incubations of *Sphagnum* peat from a temperate bog by enriching for polyphenol oxidizers and polysaccharide degraders, enhancing substrate flow toward methanogenesis under subsequent anoxic conditions. To assess whether shifts in soil microbiome structure and function operate similarly across wetland types, here we examined the sensitivity of different wetland soils to transient oxygenation. In slurry incubations of *Sphagnum* peat from a minerotrophic fen, and sediments from a freshwater marsh and saltmarsh, we examined temporal shifts in microbiomes coupled with geochemical characterization of slurries and incubation headspaces. Oxygenation did not affect microbiome structure and anoxic CH_4_ production in mineral-rich fen-origin peat and freshwater marsh soils. Key taxa linked to O_2_-stimulated CH_4_ production in the bog-origin peat were notably rare in the fen-origin peat, supporting microbiome structure as a primary determinant of wetland response to O_2_ shifts. In contrast to freshwater wetland experiments, saltmarsh geochemistry—particularly pH—and microbiome structure were persistently and significantly altered post-oxygenation, albeit with no significant impact on greenhouse gas emissions. These divergent responses suggest wetlands may be differentially resilient to O_2_ fluctuations. With climate change driving greater O_2_ variability in wetlands, our results inform mechanisms of wetland resiliency and highlight microbiome structure as a potential resiliency biomarker.

## INTRODUCTION

Wetlands are globally important ecosystems that store ∼30% of the terrestrial soil carbon pool (1), despite covering only 5-8% of the land surface (2). Their ability to act as massive carbon sinks stems from the decoupling of primary production and respiration – the typically anoxic waterlogged conditions diminish microbial decomposition of organic matter, promoting carbon preservation relative to oxic conditions (3–5). This carbon stock is still subject to microbial activity, and as such wetlands contribute significantly to the global emission budgets of potent greenhouse gases, including carbon dioxide (CO_2_), nitrous oxide (N_2_O) and in particular, methane (CH_4_) (6–8). The rapid increase in atmospheric CH_4_ levels observed in recent years has been attributed partly to increasing wetland CH_4_ emissions (9–11). Because of the highly diverse and dynamical nature of biogeochemical conditions, wetlands are among the largest and most uncertain sources of CH_4_ to the atmosphere (9). Therefore, accurately accounting for and predicting wetland CH_4_ emission trajectories is vital for climate change mitigation.

As wetland biogeochemistry is tightly linked to water table depth (12–14), changes in the frequency and duration of dry-wet cycles and associated oxygen (O_2_) and redox shifts will likely cause major changes to wetland biogeochemical cycling (14–16). Anoxic, reducing conditions that prevail during high water table periods can be conducive for methanogenesis (17). In contrast, drops in the water table resulting from droughts or decreased precipitation can oxygenate soils, stimulating CH_4_ consumption by aerobic methanotrophs while simultaneously suppressing methanogenesis in oxic layers (18,19). Further complicating the picture, however, is the recent discovery of CH_4_ emissions from oxic wetland soils (20–23). This “methane paradox” has been generally attributed to anoxic microsites harboring methanogens in otherwise bulk oxic soils (22,23). Oxygen variability, thus, clearly imparts a complex yet strong control on wetland CH_4_ fluxes. Deciphering the mechanistic underpinnings of oxygen’s roles in wetland CH_4_ emissions is a key priority for tracking and predicting global CH_4_ emission trajectories across spatio-temporal scales.

Oxygen controls complex carbon degradation in wetland soils by acting as a terminal electron acceptor for aerobic respiration, and by regulating the activity of microbial exoenzymes. Microbes produce diverse classes of exoenzymes to facilitate the breakdown of complex wetland organic carbon, primarily composed of polysaccharides (e.g., cellulose, hemicellulose) and polyphenolics (e.g., lignin, tannins) (24,25). Polysaccharide hydrolysis, which does not require O_2_, generates simpler oligomers that are readily respired or fermented. In contrast, polyphenolics degradation is generally O_2_-dependent as aromatic oxidases, commonly termed “phenol oxidases”, directly use O_2_ as a co-substrate for the oxygenation reaction (26–28). Anoxic conditions, therefore, typically lead to polyphenol accumulation. However, recent genomic and geochemical evidence suggests active polyphenol breakdown even under anoxia (29,30). The magnitude of these fluxes under different conditions and their relative importance for wetland carbon budgets remain to be ascertained.

In the absence of significant removal, accumulating polyphenols can curtail polysaccharide hydrolysis by binding to hydrolase enzymes, and thus, effectively resulting in an “enzyme latch” on carbon degradation (26–28). This “enzyme latch” hypothesis was proposed as a mechanism critical for wetland carbon storage (26–28). Many studies, however, have found contrasting evidence for the existence or strength of such a latch in various wetland systems (31–33), indicating underexplored complexity in using the latch hypothesis to explain the wetland carbon fate.

Previous experiments on *Sphagnum* peat from a peat bog in Ward Reservation, MA (herein “Ward peat”) revealed that an enzyme latch likely existed in this system as oxygenation significantly enhanced anoxic CH_4_ production in wetland soil incubations (24). Geochemical and microbial data unraveled the underlying mechanisms: during the oxic period, aerobic respiration accelerated the breakdown of complex aromatic compounds, which removed the O_2_-sensitive latch on hydrolytic enzyme activity, thereby promoting the degradation of large polymeric compounds. This generated labile carbon compounds that were readily fermented during the subsequent anoxic period, enhancing substrate flux towards methanogenesis (24,34). Specific groups of microbes were identified as key mediators of the above processes, namely (i) the proteobacterial genus *Novosphingobium*, which contain O_2_-dependent aromatic compound oxygenases and were enriched during the oxic period; (2) the acidobacterial genus *Terracidiphilus* with the ability to break down complex polymers; (3) another acidobacterial genus *Holophaga*, specializing in hydrogen (H_2_)-evolving fermentation of labile carbon compounds; and (4) *Methanobacteria*, which are hydrogenotrophic methanogens (24,34). Transient O_2_ exposure of the bog *Sphagnum* peat thus led to a cascade of biogeochemical changes, eventually leading to ∼2000-fold higher CH_4_ yields compared to continuously anoxic controls (24).

Since phenolic compounds and carbohydrate polymers comprise the majority of organic matter in typical *Sphagnum* peat (35–38), processes such as transient oxygenation that can accelerate the breakdown of these compounds (29,31,32,39,40) would greatly mobilize the carbon stock and enhance CH_4_ emissions. However, extrapolating the above observations to different wetland soils is challenging due to their heterogeneous biogeochemistry and microbial dynamics (24,41–43). The observation that aerobic breakdown of phenolics with oxygenation stimulated CH_4_ emissions in *Sphagnum* peat (24) is consistent with the enzyme latch theory. However, in iron-rich soils, abiotic reactions during anoxic-oxic oscillations can also lead to phenolic degradation (e.g., 38–42). The enzymatic latch, thus, may not be universally present across wetlands due to variability in biogeochemical factors, including polyphenol levels, and microbiome structure. In addition, variations in terminal electron acceptor pools should shift substrate cascades away from terminal CH_4_ production. O_2_ variability, therefore, may have limited impact on anoxic CH_4_ emissions, depending on wetland soil type.

Considering the diverse biogeochemistry of wetlands, we sought to determine the variability in the response of different wetland soils to transient oxygenation. Addressing this question is vital for understanding CH_4_ emissions with ongoing and projected changes in the hydrological cycle that are poised to increase the frequency of dry-wet transitions (49,50). Here, we assessed the response of three different wetland soils to brief periods of oxygenation, and compared the results to previous work, in particular to one that found O_2_-stimulation of CH_4_ release in *Sphagnum* peat from a temperate bog (24); and another that did not find stimulation of CH_4_ emissions to O_2_ shifts in a peat fen (51). Our experiments examined (i) *Sphagnum* peat from a forested wetland in the New Jersey (NJ) Pine Barrens; (ii) a mineral-soil freshwater wetland in central NJ; and (iii) a saltmarsh located along the NJ coast. We hypothesized that the enzymatic latch, if present in these wetland soils, would be weakened or eliminated with O_2_ exposure, thereby amplifying anoxic CH_4_ emissions. Additionally, this effect may be dampened by the presence of alternative terminal electron acceptors for respiration, which divert substrate flow away from methanogenesis under anoxia (52–54). To uncover potential mechanisms underlying wetland soil responses to transient oxygenation, we utilized a controlled laboratory incubation methodology with coupled measurements of microbial and biogeochemical variables.

## MATERIALS AND METHODS

### Wetland sample collection, slurry preparation, and incubation set up

Three biogeochemically distinct wetland systems were chosen for sample collection: a *Sphagnum* moss-dominated minerotrophic fen in the New Jersey Pine Barrens, a freshwater marsh in the Charles Rogers Wildlife Refuge in Princeton, NJ, and a saltmarsh in the Swan Point State Natural Area in southern New Jersey coast. At each location, wetland soil was collected from the top ∼10 cm beneath the litter/moss layer, if present. In the Pine Barrens, the *Sphagnum* peat fen was located on the banks of Bisphams Mill Creek (39.922 N, 74.591 W), in an area interspersed with Pitch Pines and various shrubs. The freshwater marsh vegetation in the Charles Rogers marshland (40.326 N, 74.659 W) was predominantly composed of cattails. Saltmarsh sediments were collected from the inundated margin of a shallow pond (40.0324 N, 74.0777 W), neighboring patches of *Spartina* grass.

At each location, acid-cleaned UV-irradiated polyethylene bags were filled with peat or sediments and transported on ice back to the laboratory. Site water (∼2 L each) was also collected from each location in acid-washed Nalgene bottles. Once in the laboratory, plant roots and litter were removed from the peat/sediments. Slurries were prepared by blending peat/sediments with 0.2 mm-filtered site water in a 1:3 volume ratio (24). The slurries were then flushed with ultrahigh purity 100% N_2_ (Airgas) for 15 minutes, and while continuing to flush with N_2_, 60 ml of each slurry was pipetted into acid-washed and autoclaved 160 ml serum bottles. The bottles were immediately closed with butyl rubber stoppers and crimp sealed.

A total of 24 slurry incubations were prepared for each wetland type. Half of these were set up as continuously anoxic controls by flushing with N_2_ gas. The remaining half were flushed with Breathing Quality air (21.5 ± 2% O_2_ and 78.5 ± 2% N_2_; Airgas). After flushing, the bottles were incubated at 20 C, shaking on their side at 150 rpm in the dark. During the oxic period of the incubation, half of the microcosms were kept aerobic by flushing the bottles with Breathing Quality air after every two days. During each flush, the continuously anoxic controls were flushed with 100% N_2_ gas. In the long-term incubation experiment, oxic conditions were maintained in the O_2_-shifted microcosms for a week, after which all bottles were flushed with 100% N_2_, thus establishing anoxic conditions in all. The subsequent anoxic incubations lasted 358 days, during which we tracked differences in biogeochemistry between the treatments. This involved monitoring of headspace gas compositions as well as destructive sampling of triplicate incubations at four timepoints (i.e., days 7, 21, 229 and 365). In a follow-up short-term incubation experiment with the Pine Barrens peat, the O_2_-shifted samples were kept aerobic for a month and were made anoxic afterward, with the anoxia lasting 65 days.

### Slurry sampling and geochemical measurements

Headspace gas samples were collected periodically to monitor CO_2_, CH_4_, and H_2_ concentrations. Using a sterile Luer lock syringe (BD), 15 ml of the headspace gas sample was collected and stored in pre-evacuated, crimp-sealed amber serum vials. Gas volumes removed from the incubation headspaces were balanced by adding either 100% N_2_ or air containing 20% O_2_, as appropriate. Final gas concentrations were adjusted for headspace dilution. Concentrations of CH_4_, CO_2_, and H_2_ in the headspace samples were analyzed using a Shimadzu GC-8A gas chromatograph equipped with a Restek ShinCarbon ST column and thermal conductivity detector. Standard curves were prepared using a calibration mixture containing 1% by volume of each target species (Airgas).

On days 7 (end of the oxic period), 21, 229, and 365, three microcosms per treatment were destructively sampled. Slurry samples for DNA/RNA (1 ml each) were aliquoted into duplicate sterile 2 ml cryotubes and stored at −80 C until extractions. 100 mL each of the slurry was used for determining ferrous ion concentrations. Another 1 ml each was used for measuring dry weight and organic matter content. Remaining slurry was centrifuged for 20 minutes at 10,000 rpm to separate out the solid and liquid fractions. The supernatant was 0.2 μm-filtered and aliquoted into clean serum vials (acid washed and combusted) for dissolved organic carbon, volatile fatty acids (VFAs), and phenolic content analyses. Solid fractions were saved at −20 C for nutrient analyses (total carbon, total nitrogen, and anions).

Slurry pH was determined by using a waterproof pH meter (Oakton) and immersing the electrode in 1 ml of the slurry sample aliquoted to a sterile conical tube (Corning). Dry weight was determined by oven-drying the slurry samples in ceramic crucibles at 105 C to a constant weight. Percent organic matter was estimated using the loss-on-ignition method, by combusting dried slurry samples at 505 C for 3 hours. Ferrous ion and total iron concentrations were measured using the Ferrozine colorimetric assay.

Standard curves were prepared by dissolving ferrous sulfate in 0.5 N hydrochloric acid. Water-extractable phenolics in the slurry supernatant was measured using the Folin-Ciocalteu colorimetric assay, using a standard curve prepared using gallic acid. Concentrations were calculated as mg/ml gallic acid equivalent. Acetate concentrations were measured on an Agilent HPLC equipped with Bio-Rad Aminex HPX-87H (300 mm × 7. 8 mm) and a DAD detector.

Infrared spectra of filtered supernatants were collected on a Bruker VERTEX 80v spectrometer equipped with a liquid-nitrogen-cooled mercury cadmium telluride (MCT) detector. Aliquots of 1 μL solution were added to the surface of a diamond ATR (attenuated total reflectance) crystal mounted on a Golden Gate ATR sample stage (Specac), and the solution was dried under moisture- and CO_2_-free air (Spectra 30 FTIR purge gas generator, Parker Balston). This step was repeated 5 times until a thin film was formed. The sample spectrum was measured as the average of 2000 scans at a scan velocity of 80 kHz with a 5 mm aperture setting. A blank spectrum was collected immediately before each sample spectrum and subtracted. All spectra were collected at 4 cm^−1^ resolution. Baseline corrections were performed on the ATR spectra using the OPUS software (Bruker). IR spectra were imported to R (v4.2) and analyzed using the package ChemoSpec (v6.1.10; (55)).

All statistical analyses and plotting of the geochemical datasets were carried out in R (v4.2), using various packages, including stats (v4.4.1; (56)) and tidyverse (v2.0.0; (57)) core packages and add-ons.

### Nucleic acid extractions, 16S rRNA gene sequencing, and microbiome analyses

Total RNA and DNA were co-extracted from the slurry samples (Table S1) by combining the RNeasy PowerSoil Total RNA kit with the RNeasy PowerSoil DNA Elution kit (Qiagen), using manufacturer’s protocols. Nucleic acid concentrations were measured using Qubit 4 Fluorometer (Thermo Fisher), and their quality was assessed using a NanoDrop spectrophotometer (Thermo Fisher). Samples were stored at −80C for later analyses.

The V4 region of the 16S rRNA gene was sequenced using Illumina MiSeq at the Princeton Genomics Core facility. Demultiplexed raw reads were primer trimmed using Cutadapt (v1.18; (58), and then, quality filtered, denoised, and merged with DADA2 (v1.26; (59)) in R (v4.2.1; (56)) to identify Amplicon Sequence Variants (ASVs). ASVs were assigned taxonomy using the Silva database (r138.1; (60)). DADA2 outputs, including taxonomic assignments and ASV read counts, were imported into PhyloSeq (v1.38; (61)) for further analyses (all performed in R v4.1.2). Eukaryotic, including mitochondrial and chloroplast, sequences were filtered out. During pre-processing, rarefaction curves were calculated separately for each wetland type to determine outliers to discard from analyses. As a result, samples without a plateauing rarefaction curve were discarded. The final pre-processed dataset included 28 Pine Barrens peat samples, 22 freshwater marsh samples, and 20 saltmarsh samples, including several technical replicates. All analyses were performed separately for each wetland type. Alpha diversity (Chao1 and Inverse Simpson) was measured using the plot_richness function in Phyloseq. The tax_fix function in microViz (v0.10.2; (62)) was used to fix missing values in the taxonomy tables. Bar plots of relative abundances were then obtained using the microshades (v1.10; (63)) package. Differential abundance testing was performed at the genus level by using Analysis of Composition of Microbiomes with Bias Correction (64,65) as implemented in the ANCOMBC package (v1.4).

### Metagenome and metatranscriptome sequencing and analyses

Nucleic acid samples passing the required quantity and quality thresholds (minimum yield of 0.5 mg and acceptable purity ratios) were used for metagenomic and metatranscriptomic sequencing. For each wetland type, one of the three replicates for each treatment across timepoints were selected for metagenome sequencing. This amounted to 9 metagenomes each per wetland type (i.e., initial T0 sample, and 4 timepoints each for the O_2_-shifted and continuously anoxic treatments). The timepoints correspond to days 0, 7 (end of the oxic period), 21, 229, and 365. Since the saltmarsh samples from later timepoints did not yield good quality DNA (Table S1), only 5 metagenomes were obtained for these, corresponding to time zero, and 2 timepoints each for the O_2_-shifted and continuously anoxic incubations. Metatranscriptomes were obtained for several of the PB peat and saltmarsh incubations, spanning two timepoints per treatment.

Paired-end sequences were obtained using the Illumina NovaSeq platform (S1 300nt flowcell) at the Princeton Genomic Core facility. Demultiplexed metagenomic reads were quality filtered using FastQC (v0.11.9; (66)) and Trimmomatic (v0.39; (67)). Anvi’o (v7.1; (68)) was used for assembly, mapping, and contig functional assignments. Metagenomes for each wetland type were co-assembled using MEGAHIT (--min-contig-len 1000; v1.2.9; (69,70)). Bowtie 2 (v2.3.5; (71)) was used to map reads to the assembled contigs. Contig functional annotations were obtained by using the functions “anvi-run-hmms” and “anvi-run-ncbi-cogs”. Demultiplexed metatranscriptomes were quality filtered using FastQC (v0.11.9; (66)) and cutadapt (v2.10; (58)). QC-filtered reads were mapped to co-assembled metagenomic contigs for each wetland type. Mapping was performed using bowtie2 (v2.3.5; (71)) in Anvi’o (v7.1; (68)). Metagenome contigs were binned using MetaBAT 2 (v1.12.1; (72)), MaxBin 2 (v2.2.7; (73)), and CONCOCT (v1.1.0; (74)). Bin refinement was carried out using the bin_refinement module in MetaWRAP (v1.2; (75)). Prodigal (v2.6.3; (76)) was used to obtain amino acid sequences of gene calls for each bin. These were uploaded to the GhostKoala server (77) to obtain KO annotations. Refined bins along with the corresponding KO annotations were exported into the merged Anvi’o profile as a collection. Relative distributions of assembled genomes and predicted functions across the metagenomes and metatranscriptomes were estimated using the “anvi-summarize” function. Additional functional annotations for the assembled genomes were obtained via DRAM (78) and RASTtk (79) within the KBase platform (80).

## RESULTS AND DISCUSSION

### Fen-origin PB peat geochemistry resilient to the O_2_ shift

#### Headspace gases

Transient O_2_ exposure did not stimulate CH_4_ emissions in fen-origin *Sphagnum* peat from the New Jersey Pine Barrens (herein “PB peat”), in contrast to the bog-derived peat incubations (herein designed “Ward peat”) (24), highlighting heterogeneous responses of peat types to O_2_ shifts (Fig. 1a,b vs. c). Regardless of the duration of the oxic period (1 week versus 4 weeks), there was no significant difference in CH_4_ production between O_2_-shifted samples and anoxic controls (Fig. 1a, S1). Like in Ward peat, oxygenation delayed the onset of methanogenesis: CH_4_ was detected in the O_2_-shifted incubation headspaces only after ∼4 weeks of anoxia following the oxic period (Fig. 1a). In comparison, within a week of anoxic incubations, detectable CH_4_ was present in the continuously anoxic control incubations (Fig. 1a). As the experiment progressed, overall CH_4_ yields were similar between the O_2_-shifted and continuously anoxic PB peat incubations (Fig. 1a), underscoring the absence of any O_2_-stimulated enhancement of methanogenesis. In comparison, the O_2_-shifted Ward peat incubations generated up to >1000-fold higher amounts of CH_4_ compared to continuously anoxic controls (Fig. 1b), clearly demonstrating a key role for O_2_ in peat carbon degradation under anoxic conditions in this system.

**Figure 1:**
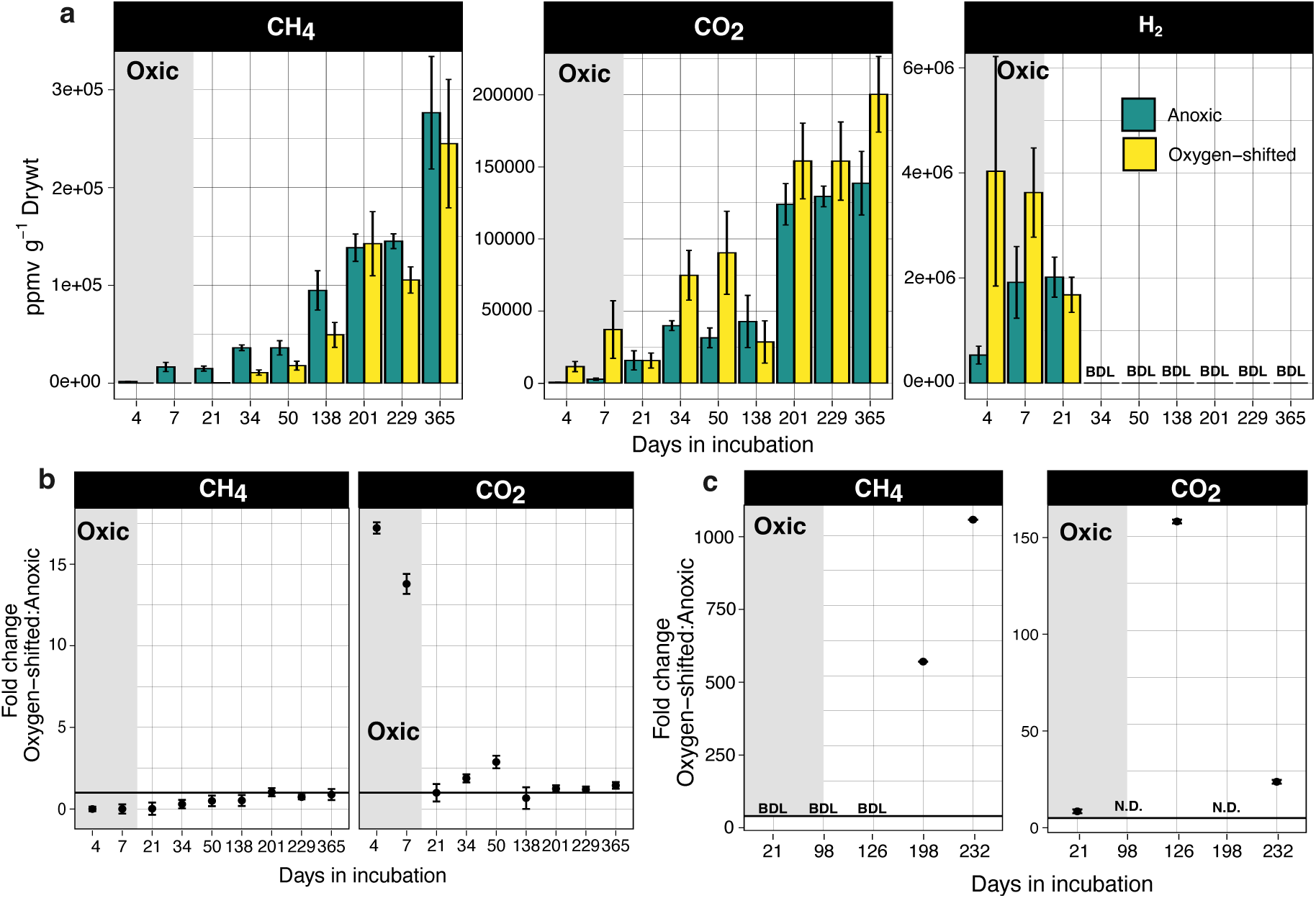
Trace gases measured in the fen-origin PB peat incubation headspaces (a,b) and compared to those in bog-origin Ward peat (c). **(a)** For PB peat, concentrations of CH_4_, CO_2_, and H_2_ across incubation time points. The grey rectangle indicates the oxic period for the O_2_- shifted samples. Error bars are standard errors around the mean of three replicates. **(b)** For PB peat, fold change in headspace CH_4_ and CO_2_ concentrations between the O_2_-shifted and continuously anoxic incubations. A fold change of 1 indicates no difference between the two treatments. (c) For Ward peat, fold change in CH_4_ and CO_2_ concentrations between O_2_-shifted and continuously anoxic incubations. Data replotted from ref. 24. Oxic period is indicated by the grey rectangle in each plot. N.D., not determined. BDL: below detection limit.

O_2_ exposition also led to higher anoxic CO_2_ emissions in Ward peat, but not in PB peat (Fig. 1b, 1c). In the latter, a sharp increase in CO_2_ emissions was observed initially upon oxygenation (Fig. 1a, 1b), indicating aerobic respiratory breakdown of peat organic carbon. The O_2_-shifted PB peat incubations yielded ∼15-fold higher amounts of CO_2_ during the oxic period compared to the anoxic controls (Fig. 1b). Following the onset of anoxia, the two treatments were largely similar in CO_2_ emissions, indicating that any O_2_-induced change in CO_2_-forming metabolism was short-lived (Fig. 1a, 1b). This was in stark contrast to the Ward peat, which showed much higher (∼20- to 150-fold) CO_2_ emissions in the O_2_-shifted incubations compared to anoxic controls long into the anoxic period following the O_2_ shift (Fig. 1c).

These observations indicate that the brief O_2_ exposure enhanced CO_2_-forming carbon mineralization in Ward peat in addition to promoting methanogenesis, likely by increasing the flux and availability of labile compounds for microbial degradation which persisted into the anoxic period. Such an effect appears to be absent in the PB peat, where O_2_-mediated CO_2_-forming respiration during the oxic period likely did not yield a similar influx of labile C for further anoxic degradation.

The headspace hydrogen (H_2_) evolution trajectories provide further evidence for the lack of O_2_ stimulation of anoxic metabolism (specifically fermentation) in PB peat compared to Ward peat. Both the O_2_-shifted and continuously anoxic PB peat incubations showed high levels of H_2_ in the headspace during the first three weeks of incubation (Fig. 1a). H_2_ levels dropped below the detection limit by week 3 and remained undetectable for the remainder of the experiment (Fig. 1a). In contrast, significant amounts of H_2_ were detected in the Ward peat incubation headspaces even at the final incubation timepoint (day 232, 134 days anoxic; Fig. S2; (24)). In addition, acetate, another key product of microbial fermentation, accumulated at millimolar levels in the Ward peat (up to ∼2.5 mM in O_2_-shifted Ward peat and ∼7 mM in the anoxic controls; (24). In contrast, no measurable acetate was detected in the PB peat incubations at any timepoint during the incubation (detection limit 560 mM).

#### Aqueous phase geochemistry

O_2_ exposure further led to a decline in pH and Fe(II) concentrations in the PB peat incubations, which coincided with a sharp increase in sulfate (SO_4_^2-^) levels (Fig. 2), likely derived from the oxidation of reduced organosulfur compounds that can be abundant in peat (81) or iron-sulfides (82,83). Fe(II) and SO_4_^2-^ levels were reestablished to prior values upon making the incubations anoxic (Fig. 2), potentially indicating the coupled effect of microbial transformations such as sulfate reduction and the regeneration of reduced iron- and sulfur-compounds. At least at the very beginning of the anoxic period in the O_2_- shifted peat, SO_4_^2-^ and Fe(III) may have served as alternative electron acceptors for microbes. However, SO_4_^2-^ levels remained well-below the limiting concentration for microbial sulfate reduction (300 mM; (81,84)), and therefore, the importance of cryptic microbial sulfate reduction in PB peat remains unclear.

**Figure 2:**
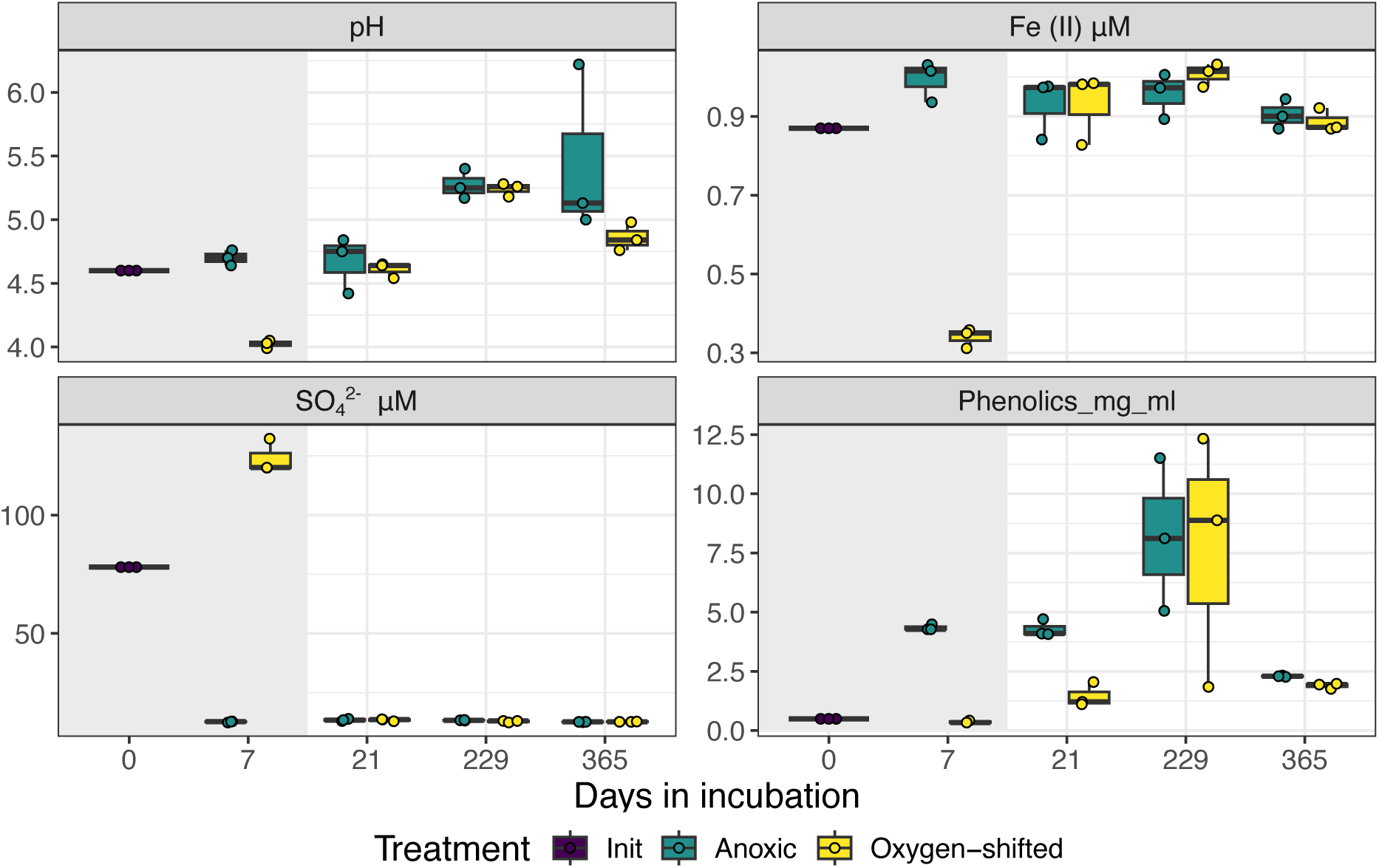
Changes in wetland soil geochemistry with the O_2_ shift. Key geochemical variables measured in PB peat across timepoints in the experiment. Oxic period is indicated by the grey rectangle. Three replicate measurements are included in each boxplot (overlayed dots depict the actual values). N.D., not determined. Phenolics concentrations were measured as mg/ml gallic acid-equivalent.

Collectively, the geochemical observations indicated carbon degradation pathways in PB peat to be largely unaffected by the O_2_ shift, likely due to a weak enzymatic latch in this peat. This is further evidenced by the lack of measurable phenolics in the initial PB peat sample (Fig. 2). Total phenolics were ∼2- to 5-fold higher in the anoxic controls compared to the O_2_-shifted samples on day 7 and 21 (Fig. 2), indicating O_2_-mediated breakdown of polyphenols in the O_2_-shifted samples. Following the oxic period, FTIR data indicated the mobilization of small organic molecules, including aromatics (C=C at 1627, 1594, and 1493 cm^-1^), amides (amides I and II bands at 1655 and 1579 cm^-1^, respectively), and aliphatics (C-H between 2700 and 3000 cm^-1^), which were absent in the anoxic controls and diminished at later timepoints, suggesting microbial consumption (Fig. S3). The anoxic and O_2_-shifted samples had largely similar IR spectra at later timepoints during anoxia (Fig. S3), suggesting a reset of soluble organic matter composition following oxygenation. This contrasts starkly with the Ward peat, where FT-ICR-MS data revealed major changes in organic matter composition, notably significant O_2_-enhanced removal of aromatic compounds (24).

Overall, geochemical measurements point to differences in the controls on microbial carbon metabolism in the two peats. Even a relatively short period of O_2_ pre-exposure was sufficient to enhance anoxic period CH_4_ release in Ward peat (e.g., ∼55-fold increase in anoxic CH_4_ emissions following a week of O_2_ exposure; (24)). The apparent absence of a similar effect in our experiments with PB peat (Fig. 1a), even with a longer period of O_2_ pre-exposure of 30 days (Fig. S1) suggests fundamentally different O_2_ controls on microbial metabolisms and carbon cycling in these two *Sphagnum* peats. In addition, the lack of accumulation of fermentation byproducts in PB peat indicates either a tight coupling of fermentation with methanogenesis or alternative anaerobic respiration of organic matter in this peat. Indeed,ncreased sulfate concentrations in PB peat upon O_2_ exposure (Fig. 2) suggest the possibility of sulfate reduction as an alternative respiratory metabolism during anoxia in this peat.

### Fen-origin PB peat microbiome largely resilient to the O_2_ shift

Microbial community structure and its response to the O_2_ shift in the fen-origin PB peat contrasted strongly with that of Ward peat. O_2_-stimulation led to significant changes in the Ward peat microbiome (24,34). The PB peat microbiome, in contrast, was largely resilient to the O_2_ shift, as the two treatments were similar in community composition throughout the experiment (Fig. 3a). As the experiment progressed, the PB microbiome shifted from a community dominated by *Proteobacteria* and *Acidobacteriota* to one dominated by *Bacteroidota* (Fig. 3a). These changes were, however, independent of the O_2_ shift, as they were consistent between anoxic and O_2_-shifted samples.

**Figure 3:**
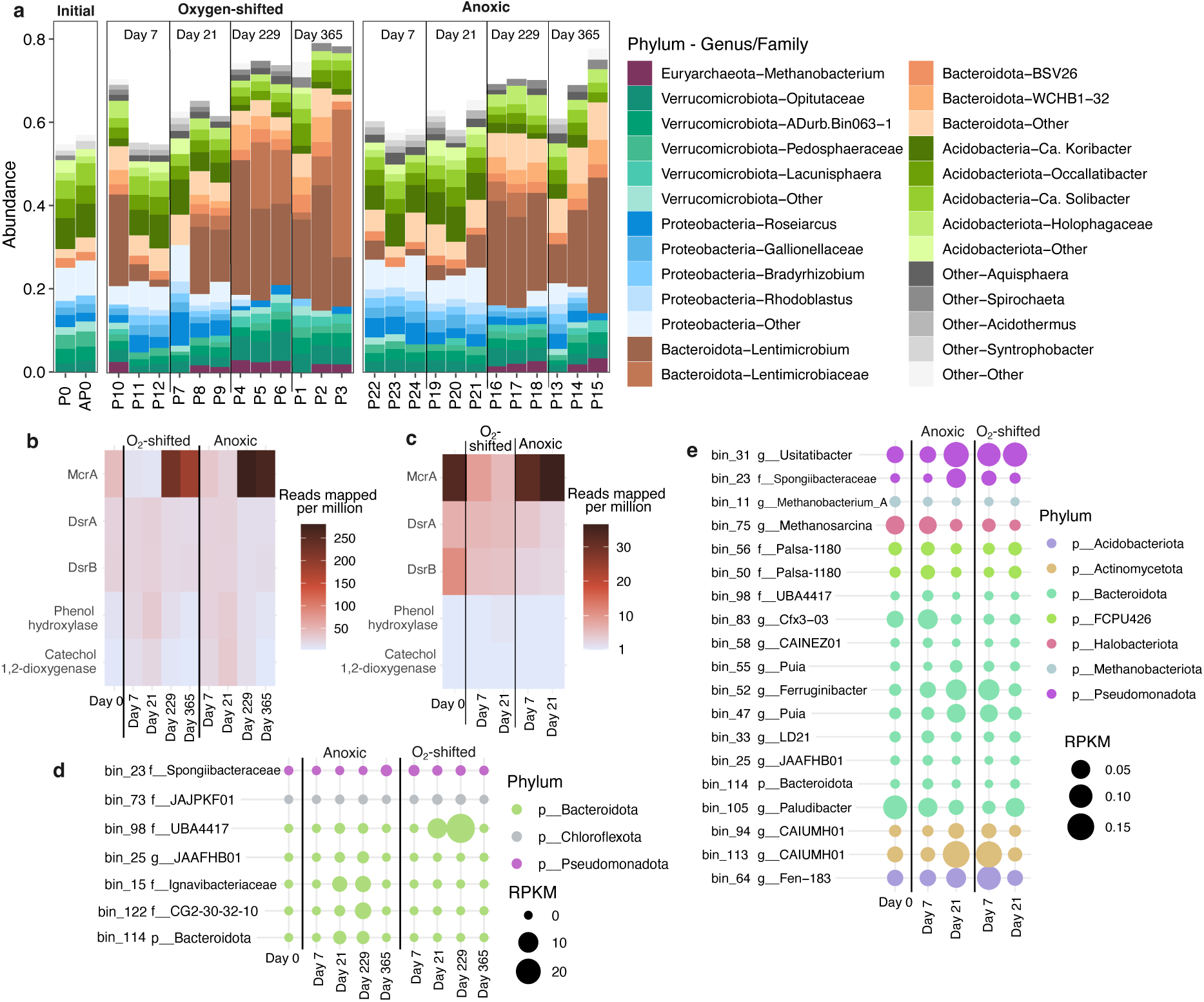
PB peat microbial community response to the O_2_ shift. Microbial community composition across incubation timepoints assessed using the V4 region of 16S rRNA gene. (**a**) Top 5 phyla comprising >50% of the community in the PB peat microbiome. Dominant genus or family-level lineages are also highlighted for each phylum. Gray box in the “Oxygen-shifted” panel indicates replicate samples at the end of the oxic period. (b) Relative abundances of selected functions in the PB peat metagenomes. McrA: methyl coenzyme reductase; DsrA and DsrB: sulfite reductase subunits A and B. (**c**) Relative abundance of the selected functions in the PB peat metatranscriptomes. MAGs identified as differentially abundant (d) and active (c) between O_2_ treatments in PB peat. For each MAG, taxonomic identity at the genus, family, or phylum-level has been presented. RPKM: reads mapped per kilobase of genomes per million total reads.

### Notable taxa and functions associated with a weaker “latch” in PB peat

Key taxa observed to be vital players in the O_2_-stimulated CH_4_ emissions in Ward peat were noticeably rare in PB peat. In Ward peat, three major microbial groups were identified as key players in the O_2_ response: *Novosphingobium*, an alphaprotebacterial genus harboring aromatic oxidases, Acidobacteria (specifically, the genera *Holophaga* and *Terracidiphilus*) capable of polysaccharide hydrolysis and fermentation, and *Methanobacterium*, a methanogenic genus of the phylum Euryarchaeota (24). *Novosphingobium* were significantly enriched in the O_2_-shifted Ward peat, comprising up to 80% of the 16S rRNA gene amplicons during the oxic period. Similarly, *Acidobacteria* were enriched under anoxia, comprising >50% of the community (24,34). These taxa were relatively rare (overall abundance <5% of the community) in the PB peat microbiome (Fig. 3a, S4). The relative abundance of *Novosphingobium* indeed increased in PB peat immediately following the O_2_ exposure, however, they still only comprised <2% of the total community (Fig. S4). *Holophaga* were present throughout the incubation period in PB peat although they constituted a very small proportion of the community (i.e., up to 5% relative abundance in PB peat versus ∼50% in Ward peat; Fig. S4). Also, unlike in the Ward peat incubations, *Holophaga* abundances did not vary between O_2_ treatments in PB peat. Similarly, the primary methanogen in the community, *Methanobacterium*, also did not increase in abundance during anoxia in O_2_-shifted PB peat unlike in Ward peat (Fig. 3a).

While O_2_ exposure did not alter the PB peat microbiome as drastically as it did the Ward peat microbiome, subtle yet short-lived compositional changes still followed the oxygenation period. There was a sharp increase in community alpha diversity with O_2_ exposure (i.e., measured as the number of observed amplicon sequence variants or ASVs; Fig. S5), as diverse microbes likely exploited the new niche for aerobic respiration. This effect was relatively short-lived as the O_2_-shifted samples progressively became less diverse over time, and significantly lower than anoxic controls at later timepoints (Fig. S5; Wilcoxon Rank Sum test with FDR adjusted p-values < 2e-16 for all). In comparison, the continuously anoxic controls showed largely invariable ASV richness throughout the experiment (Fig. S5). ANCOM differential abundance testing uncovered 24 ASVs in the PB peat microbiome as differentially abundant between the O_2_-shifted and continuously anoxic treatments (q-value < 0.001; Fig. S6). However, most of these ASVs had log Fold Change values < 1 (with the maximum being 1.5; Fig. S6), suggesting that the abundance differentials were rather small. Both treatments showed a reduction in community evenness over time (as measured using the Inverse Simpson index), suggesting an increasingly uneven distribution of microbial populations among the ASVs (Fig. S5). In other words, while the overall richness of the communities decreased over time, certain ASVs (i.e., “species”-level lineages) became increasingly dominant. Overall, the PB peat microbiome was largely resistant to the O_2_ shift, despite some differences in the relative abundances of ASV- (“species-”) level sub-lineages across the O_2_ treatments (Fig. S5, S6). Temporal changes in community composition in the PB peat microbiome (Fig. 3a) are thus, likely more reflective of the “bottle effect”, which results from changes in the quality and quantity of metabolic substrates over time. These patterns contrast sharply with Ward peat, where the microbiome composition varied significantly not only across timepoints but also between the O_2_ treatments (24,34).

Consistent with the microbiome composition data, comparisons of key functional profiles between the O_2_ treatments also highlighted the difference between the two peats in their response to transient O_2_ exposure: aromatic oxygenases (phenol hydroxylases and catechol dioxygenases) found to be enriched during the oxic period in Ward peat (24) were of notably low abundance in the PB peat metagenomes and metatranscriptomes and did not respond to the O_2_ shift (Fig. 3b, 3c). This agrees with the FTIR data showing minimal changes in aromatic compound transformations post oxygenation and a modest accumulation of likely oxidized aromatics in the aqueous phase following oxygenation (Fig. S3). Indicating potential importance of alternative electron acceptors in PB peat, sulfite reductase (DsrAB) transcripts were relatively more enriched in the O_2_-shifted PB peat than in the anoxic controls (Fig. 3b, 3c), which coincided with the higher levels of sulfate measured in the O_2_-shifted samples at the end of the oxic period (day 7; Fig. 2). Methanogenesis, as expected, was more enriched in the continuously anoxic controls compared to the O_2_-shifted samples (using the marker gene methyl coenzyme M reductase, *mcrA*; Fig. 3b, 3c).

Assembling metagenome contigs into draft genomes (i.e., MAGs) allowed a more nuanced look at the differentially abundant and active lineages across treatments and timepoints in PB peat. We obtained 72 medium- to high-quality MAGs (i.e., > 70% complete; <10% contamination) from the metagenomes (Table S2). DESeq differential abundance testing revealed 7 MAGs as being differentially abundant between O_2_-shifted and anoxic PB peat (Fig. 3d), and 19 as differentially active across treatments (i.e., differential read recruitment in the metatranscriptomes; Fig. 3e).

Both the metagenome and metatranscriptome abundance profiles indicated that lineages within major phyla respond differentially to the O_2_ shift, suggesting diverging niche adaptations and metabolic strategies among related organisms. Among the six *Bacteroidota* MAGs identified as differentially abundant between treatments in the metagenomes, only one was differentially enriched in O_2_-shifted samples while the remaining five were relatively more abundant in the anoxic controls (Fig. 3d). Functional comparisons did not reveal any discernable metabolic features between these MAGs. A *Pseudomonadota* MAG bin_23 (novel genus within family *Spongiibacteraceae*) was relatively more abundant in both the PB peat metagenomes and metatranscriptomes during the oxic period (day 7; Fig. 3d,e). In addition to the aerobic respiration machinery, this MAG harbored pathways for O_2_-dependent degradation of aromatic compounds. Among the differentially active MAGs, although most appear to be facultatively anaerobic heterotrophs, a few stood out in terms of alternative respiration capabilities. Specifically, bin_31 (*Pseudomonadota* genus *Usitatibacter*) and bin_64 (*Acidobacteriota* genus Fen-183), which were differentially abundant in the O_2_-shifted PB peat metatranscriptomes (Fig. 3e), have the potential for sulfate reduction. Bin_31 additionally has the potential for iron oxidation (Table S3). Their heightened activity during the oxic period thus coincides with the initial decrease in Fe(II) and surge in sulfate upon oxygenation of PB peat (Fig. 2).

### Freshwater marsh and saltmarsh responses to O_2_ shift

To compare the sensitivity of different types of wetlands to O_2_ shifts, we repeated the incubation experiments with sediments collected from a freshwater marsh (herein “FW marsh”) and saltmarsh. CH_4_ was detected in the FW marsh incubation headspaces only after 3 weeks of anoxic incubation (compared to 1-week of anoxic incubation of PB peat). CH_4_ yield was generally 1 order of magnitude lower than in PB peat (Fig. 4a and 1a). CH_4_ production in the O_2_-shifted FW marsh samples was initially lower compared to the continuously anoxic controls; however, over time, similar quantities of CH_4_ were measured in both treatment headspaces (Fig. 4). Thus, similar to PB peat, transient O_2_ exposure had no significant effect on CH_4_ release during the subsequent anoxic period in FW marsh either (Fig. 4a). CO_2_ yields in the FW marsh incubations were also several folds lower compared to PB peat incubations (Fig. 4a and Fig. 1a). Prior O_2_ exposure led to slightly higher CO_2_ concentrations in the FW marsh incubations compared to anoxic controls, but the difference was increasingly insignificant over time (Fig. 4a).

**Figure 4:**
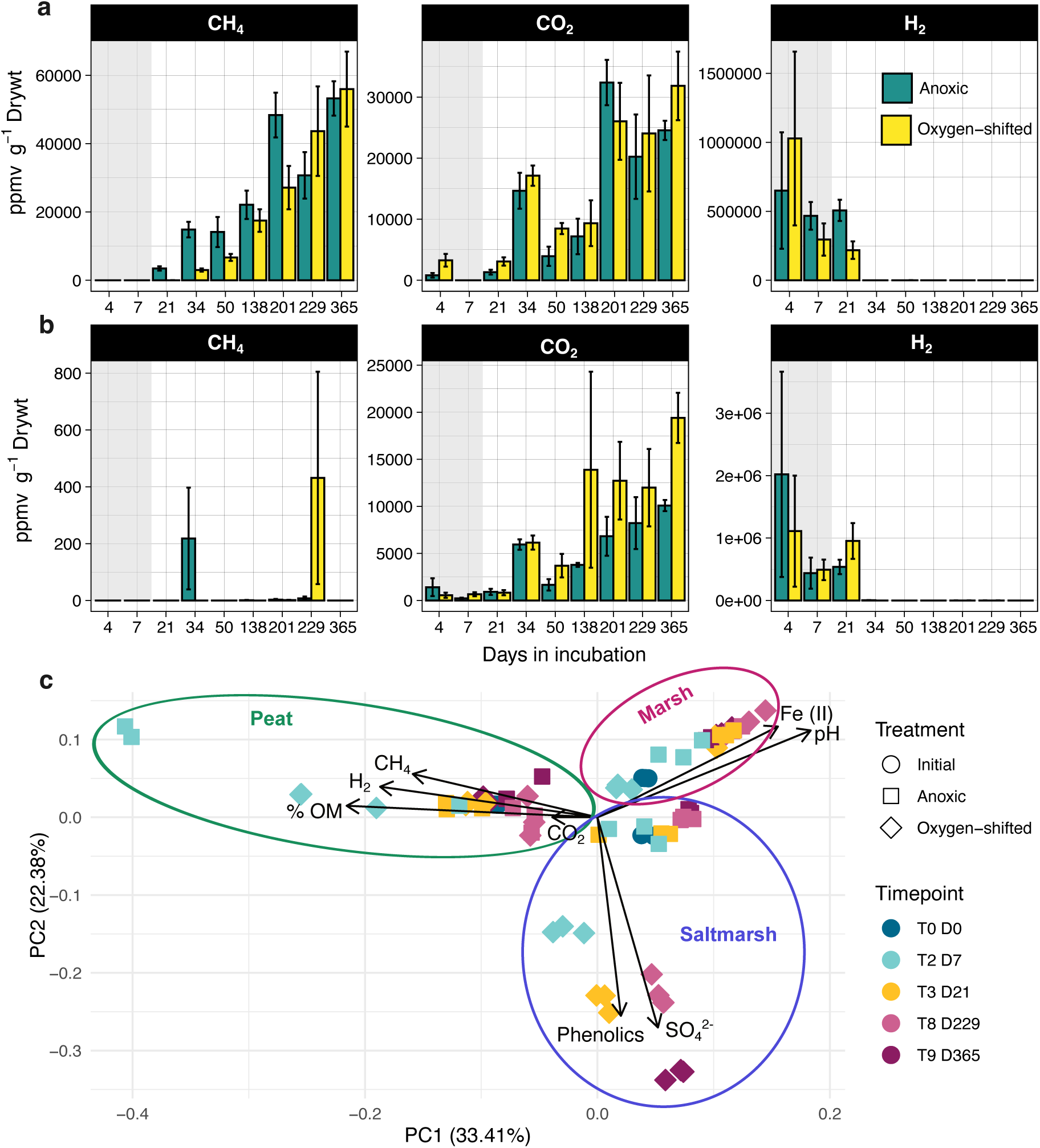
Geochemical responses of FW-marsh and saltmarsh to O_2_ shift. CH_4_, CO_2_, and H_2_ measured in the incubation headspaces across timepoints in FW marsh (**a**) and saltmarsh (**b**). The grey rectangle indicates the oxic period for the O_2_-shifted samples. Error bars are standard errors around the mean of three replicates. (**c**) PCA of key geochemical variables measured over the course of the experiment. Shapes indicate different treatments. Ellipses are drawn to indicate each wetland type.

In contrast to both PB peat and FW marsh, the O_2_-shifted saltmarsh incubations, on average, released higher amounts of CO_2_ (1.6- to 3.8-fold) during the anoxic period than continuously anoxic controls as the incubations progressed (Fig. 4b). Interestingly, this difference was only observed with long (>6 weeks) anoxic incubation of the O_2_-shifted saltmarsh samples; during and immediately following the oxic period, both treatments released similar amounts of CO_2_ (Fig. 4b). Relatively small amounts of CH_4_ were detected in the headspaces of a few of the saltmarsh incubations at a couple of timepoints during the anoxic phase (Fig. 4b); however, there appeared to have been active turnover of this CH_4_ as the levels dropped below detection at later timepoints (Fig. 4b).

Measured geochemical properties of the slurry changed quite significantly upon oxygenation in all three wetland types (Fig. 4c, S7). These changes, however, were rather short-lived in both the PB peat and FW marsh as the O_2_-shifted and continuously anoxic samples converged over time (Fig. 2, 4c, S7). Saltmarsh geochemistry, on the other hand, was more persistently altered upon O_2_ exposure: samples in the two treatments continued to diverge in their measured geochemical properties even after several months of anoxic incubations (Fig. 4c, S7). This pointed to more persistent changes in saltmarsh geochemistry upon oxygenation compared to the other two wetland types, likely driven by the significant drop in pH (∼6.5 in the initial sample to 4 in the O_2_-shifted sample; Fig. S7), and approximately twofold increase in sulfate levels (Fig. S7).

### Microbiome responses to O_2_ shifts in the freshwater marsh and saltmarsh incubations

Microbiome response to O_2_ shift in the FW marsh was strikingly similar to that in PB peat, with both communities transitioning form Proteobacteria-dominated to Bacteroidota-dominated over time (Fig. 5a). As in the case of PB peat, these changes did not reflect any effect of the O_2_-pretreatment in FW marsh either, as O_2_-shifted and anoxic samples were compositionally similar across timepoints (Fig. 5a). These observations further align with the minimal changes in geochemistry upon oxygenation (i.e., no significant difference in trace gas emissions between the two treatments and the lack of persistent changes in sediment geochemistry; Fig. 4a, 4c).

**Figure 5:**
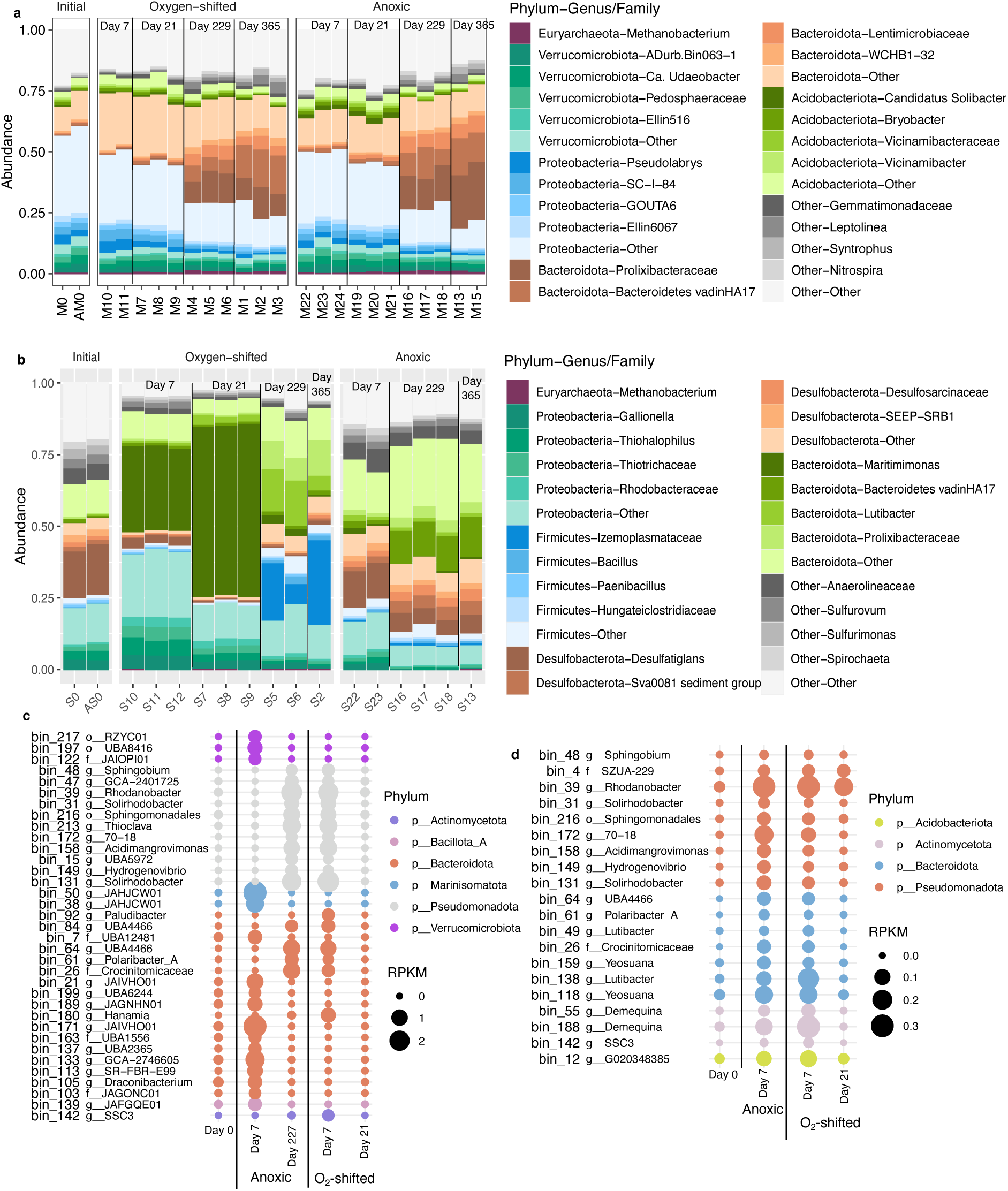
Microbiome response to O2 shift in FW-marsh and saltmarsh. Microbial community composition across time points in FW-marsh (a) and saltmarsh (b), assessed using the V4 region of 16S rRNA gene. Top 5 phyla comprising >50% of the communities are highlighted in each plot. Dominant genus or family-level lineages are also indicated for each phylum. (b) MAGs identified as differentially abundant (d) and active (c) between O_2_ treatments in the saltmarsh incubations. For each MAG, taxonomic identity at the genus, family, or phylum-level has been presented. RPKM: reads mapped per kilobase of genomes per million total reads.

The saltmarsh microbiome, in contrast, changed significantly upon O_2_-addition (Fig. 5b), as would be expected based on the notably different geochemical properties of the anoxic controls and O_2_- shifted samples (Fig. 4c). The anoxic controls were dominated by *Desulfobacterota* and *Bacteroidota* throughout the incubation period, with little compositional changes with time (Fig. 5b). The O_2_-shifted samples were differentially enriched in *Proteobacteria* and specific lineages of *Bacteroidota*, with *Firmicutes* becoming abundant towards the final timepoints (Fig. 5b). In particular, the genus *Maritimimonas* (phylum *Bacteroidota*) significantly increased in relative abundance upon oxygenation and remained dominant into early phase of the anoxic period (Fig. 5b). With prolonged anoxia, a different *Bacteroidota* genus *Lutibacter* became increasingly more abundant in the O_2_-shifted samples, along with the *Firmicutes* family *Izemoplasmataeceae* (Fig 5b). In contrast to PB peat, much higher log Fold Change values (up to 4.5 and −3.5) were observed for the differentially abundant ASVs in the saltmarsh incubations (Fig. S6c). Thus, the O_2_-shifted and continuously anoxic saltmarsh microbiomes remained compositionally dissimilar throughout the incubation period (Fig. 5b), aligning with the persistent changes in geochemistry brought about by O_2_ exposition (Fig. 4c).

Several phylogenetically novel MAGs were obtained from the saltmarsh metagenomes, including one that was classified as a new phylum-level lineage (Table S2). Aligning with the community composition patterns observed for the ASV data, many of the MAGs were conspicuously differentially abundant across the two treatments (Fig. 5c, 5d). The altered geochemistry persisting long after O_2_ exposition in the saltmarsh (Fig. 4c) likely promoted differential enrichment of microbial taxa in these incubations. Ecological or biogeochemical consequences of these observations, especially in field settings, remain to be examined.

### Broader implications

Hydrological shifts leading to O_2_ variability are a primary control on the dynamic nature of wetland greenhouse gas emissions. Given the diversity of wetlands, the specific mechanistic underpinnings of this control, particularly the effect of O_2_ shifts on microbial biogeochemistry, remain challenging to decipher, impeding our ability to accurately account for wetland contributions to climate-active trace gas emissions.

#### Microbial and geochemical factors regulating wetland response to O_2_ shifts

Several studies have reported accelerated carbon degradation (i.e., CO_2_ emissions) in response to droughts or redox shifts in wetlands (12,24,85). Such observations have typically been attributed to O_2_- stimulated breakdown of complex carbon compounds (12,24). Our results, however, highlight the heterogeneity in wetland responses to O_2_ shifts, even in peat dominated by the same plant genus, *Sphagnum*.

The stark contrast between the Ward *Sphagnum* peat and the PB forest *Sphagnum* peat reveal the variable effects of O_2_ shifts on peat CH_4_ emissions. In the Ward peat, geochemical and microbial data indicated a key role of O_2_ in promoting the degradation of complex aromatic and polymeric carbon, promoting H_2_-, acetate-, and CO_2_-evolving fermentation, and eventually leading to a surge in methanogenesis (Fig. 6). No such patterns were observed in PB peat as exemplified by several geochemical and microbial measurements, namely, (i) only minor compositional shifts in the microbiome with the O_2_ shift (Fig. 3), (ii) no enrichment of aromatic carbon degraders and associated enzymes (Fig. 3b, 3c), and (iii) the lack of accumulation of fermentation byproducts (H_2_ and acetate). While hydrogenotrophic methanogenesis seemed to be the dominant methanogenic pathway in both peats (since *Methanobacteria* were the dominant methanogens in both cases), O_2_ exposure did not result in enhanced substrate flow towards methanogenesis in PB peat unlike in Ward peat. Overall, microbial carbon degradation appears to have followed significantly different routes in the two peat types. These differences may arise from a variety of factors, including differences in the initial microbiome composition, presence of alternative electron acceptors, and the larger ecological and environmental context of each peat system.

**Figure 6:**
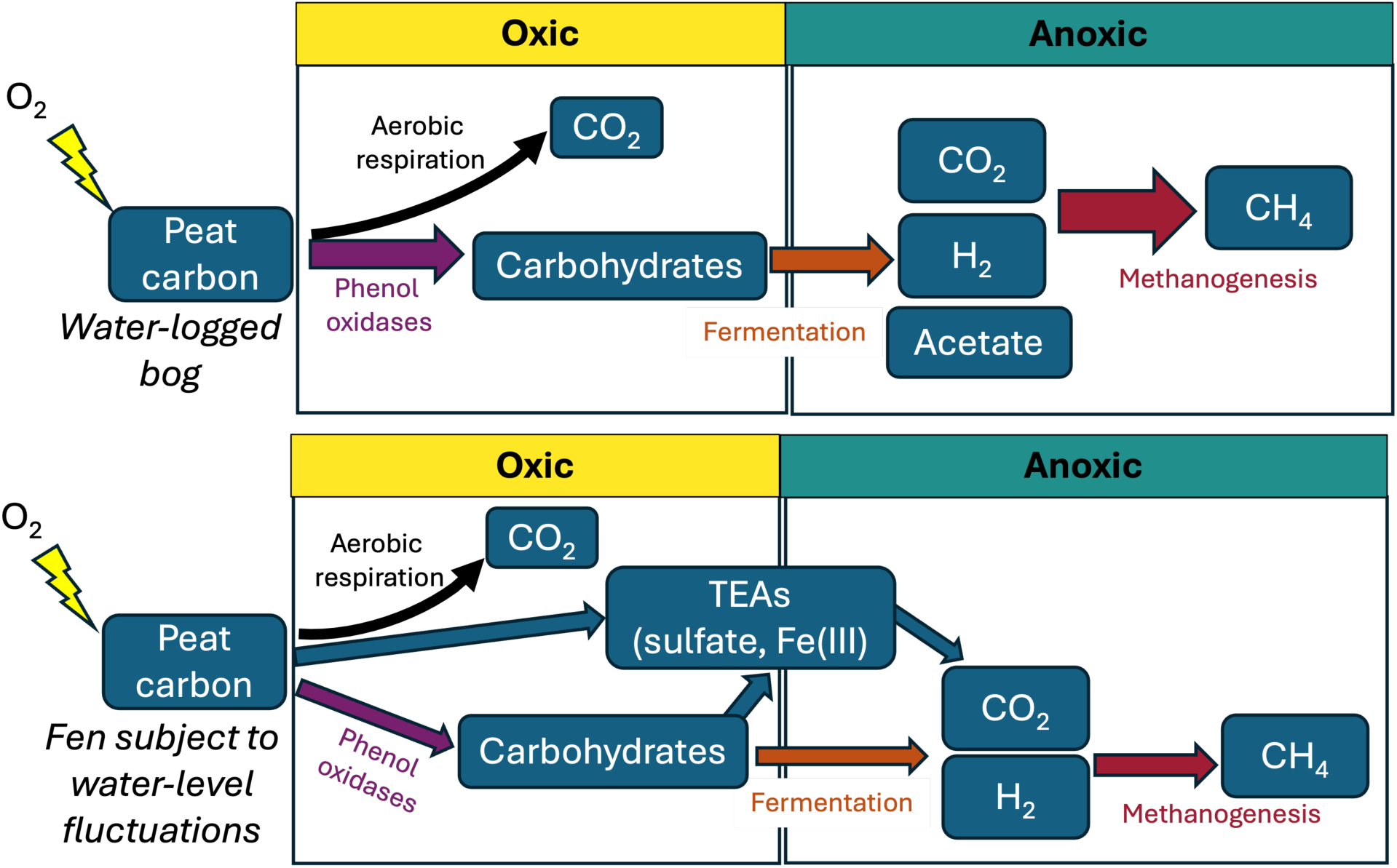
Schematic diagram of microbially-mediated carbon flow with O_2_ shift in the (a) Ward sphagnum peat compared to (b) PB Sphagnum peat. In contrast to Ward peat, transient oxygenation does not lead to a surge in anoxic CH_4_ production in PB peat, likely due to (i) the PB fen being subject to fluctuations in water (and thus O_2_) levels, favoring a microbiome with lower sensitivity to O_2_ variability, and (ii) alternative terminal electron acceptors redirecting carbon flow away from CH_4_. The width of the arrows presents the importance of relevant metabolisms.

The environmental context of each peat system potentially exerts a major control on CH_4_ biogeochemistry: while the Ward peat originated from a peat bog, the PB peat originated from a *Sphagnum*-dominated minerotrophic fen located along the banks of a freshwater creek. Given its proximity to the creek, the PB peat system likely experiences more frequent changes in water table levels compared to the Ward peat bog. This likely imparts a higher degree of tolerance for hydrological and O_2_ variability in the PB peat. These results are comparable to the observations in (51), where dry-wet manipulations in a Boreal minerotrophic fen did not stimulate phenol oxidase activity or enhance CH_4_ production, despite an initial surge in CO_2_ production. This study also reported an increase in sulfate, Fe(III), and nitrate concentrations upon soil drying. While nitrate levels in the PB peat was below detection limit throughout, the increase in sulfate concentrations we observed (∼112 uM) were largely comparable to those reported in (51) (>100 uM). At least part of the carbon flow in PB peat may have thus been directed towards anaerobic respiration, specifically sulfate reduction, which is further supported by the relatively higher expression of Dsr genes and putative sulfate reducers in the PB peat metatranscriptomes (Fig. 3b,c). These results also compare to the recent observation that carbon decomposition in Northern peatlands is likely dominated by TEA respiration rather than methanogenesis (54). Geochemical and microbial data thus indicate that unlike in Ward peat, phenolic compounds may not comprise a strong “latch” on PB peat carbon degradation. In agreement with field-based studies (51), our observations collectively suggest that O_2_ fluctuations in minerotrophic fen systems are unlikely to stimulate CH_4_ emissions.

Additionally, the strikingly dissimilar response of the saltmarsh sediments to transient oxygenation points to varying microbial community compositions and ionic interactions shaped by salinity and sulfate levels, which may cause divergent responses of brackish coastal wetlands to O_2_ shifts. Systems subjected to tidal flushing and associated periodic changes in O_2_ levels may, however, be more buffered against O_2_ variability.

#### Wetland resiliency to O_2_ shifts in the context of global change

The large heterogeneity in susceptibility of wetland carbon stock to oxygen variability (or dry-wet cycles) is well-documented (reviewed in (86)) but poorly understood mechanistically. Our observations suggesting wetland resiliency to O_2_ shifts varies with wetland type has important implications for wetland management under climate change scenarios. Compared to bogs, mineral soil wetlands such as the FW marsh, and minerotrophic peatlands supporting alternative respiration, such as PB peat, are more resilient to O_2_ shifts. Partly, this resilience could stem from “memory effects” due to frequent redox changes that could confer microbial communities with the capacity to adapt. Such resiliency may be absent in permanently anoxic sediments of saltmarshes as we observed a clear lack of convergence between the O_2_-shifted and anoxic saltmarsh samples. While the validity of these results to field settings remains to be assessed, our observations indicate potentially major, persistent changes in saltmarsh ecosystems resulting from redox variability associated with coastal restoration or sea-level rise.

Our observations inform the mechanisms underlying heterogeneous responses of wetland carbon stock to oxygen variability. Most notably, they point out specific microbial taxa as potential indicators of wetland resiliency under hydrological variations and indicate that microbiome data can inform predictions of wetland biogeochemical behavior upon redox shifts. The results also underscore the need to assess wetland resiliency in the context of their divergent ecological settings. Such careful characterization of the environmental heterogeneity is essential for accurately scaling up laboratory observations to predictive global models of peatland methane emission trajectories.

## DATA AVAILABILITY STATEMENT

All sequence data has been deposited in the NCBI Sequence Read Archive under the BioProject accession PRJNA1123272 (data will be publicly available upon publication). All other datasets and R codes used for analyses and for generating the figures are available at the GitHub repository

(https://github.com/Linta-Reji/Reji2024_wetland_O2shifts), with large files uploaded to Zenodo (doi: 10.5281/zenodo.14510955).

## CONFLICT OF INTERESTS

The authors declare no competing interests.

## ACKNOWLEDGMENTS

This study was supported by the Carbon Mitigation Initiative at the High Meadows Environmental Research Institute at Princeton University. The authors thank Dr. E. Han for assistance with sample collection. The authors are pleased to acknowledge that the work reported on in this paper was substantially performed using the Princeton Research Computing resources at Princeton University which is consortium of groups led by the Princeton Institute for Computational Science and Engineering (PICSciE) and Office of Information Technology’s Research Computing. This work was supported by the Office of Science, Office of Biological and Environmental Research, of the US Department of Energy under Award Numbers DE-AC02-05CH11231, DE-AC02-06CH11357, DE-AC05-00OR22725, and DE-AC02-98CH10886, as part of the DOE Systems Biology Knowledgebase.

## Supplementary Figures

**Figure S1:**
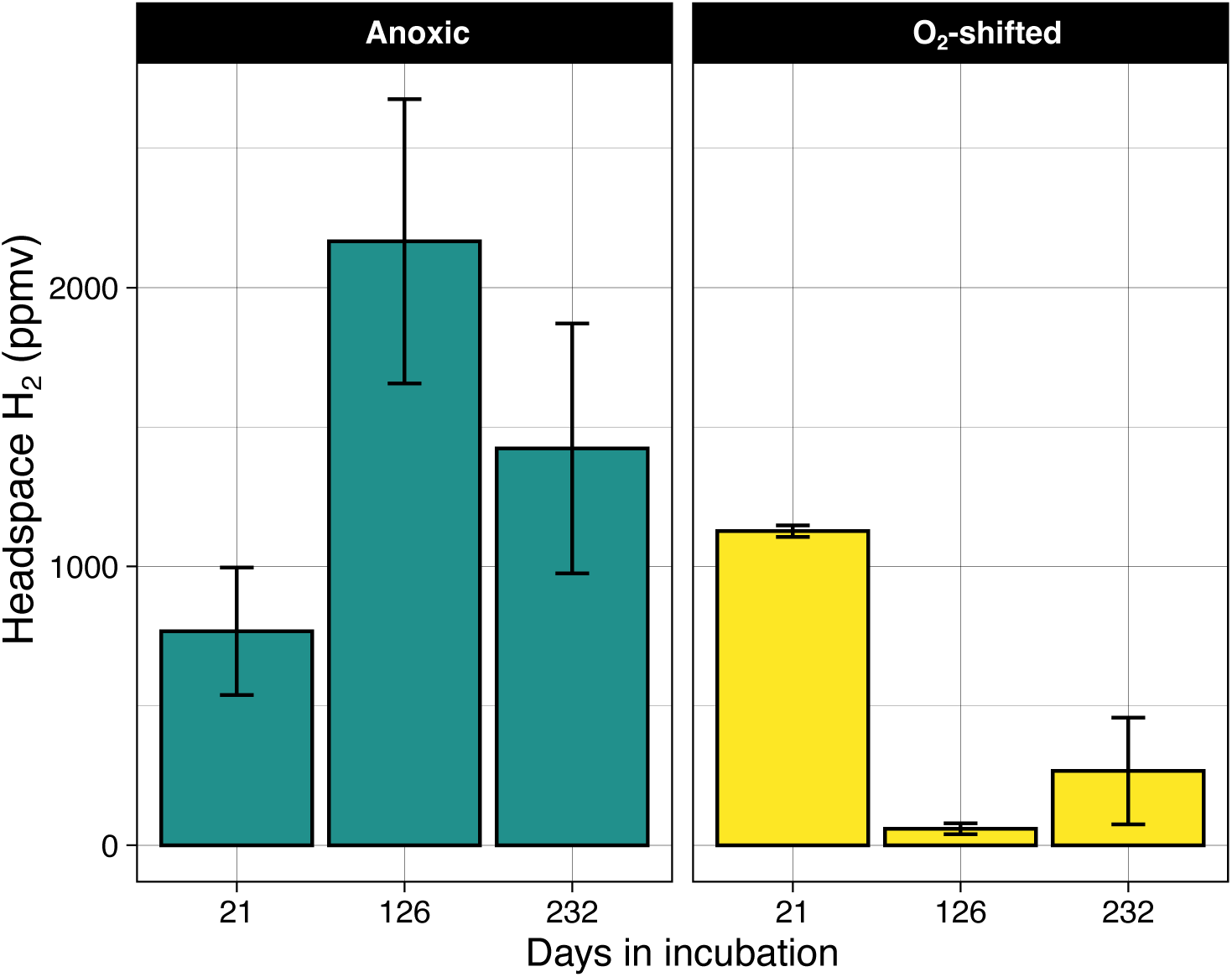
H_2_ Measured in Ward peat incubations. Error bars are standard errors around the mean of three replicates. Data re-plotted from (24).

**Figure S2:**
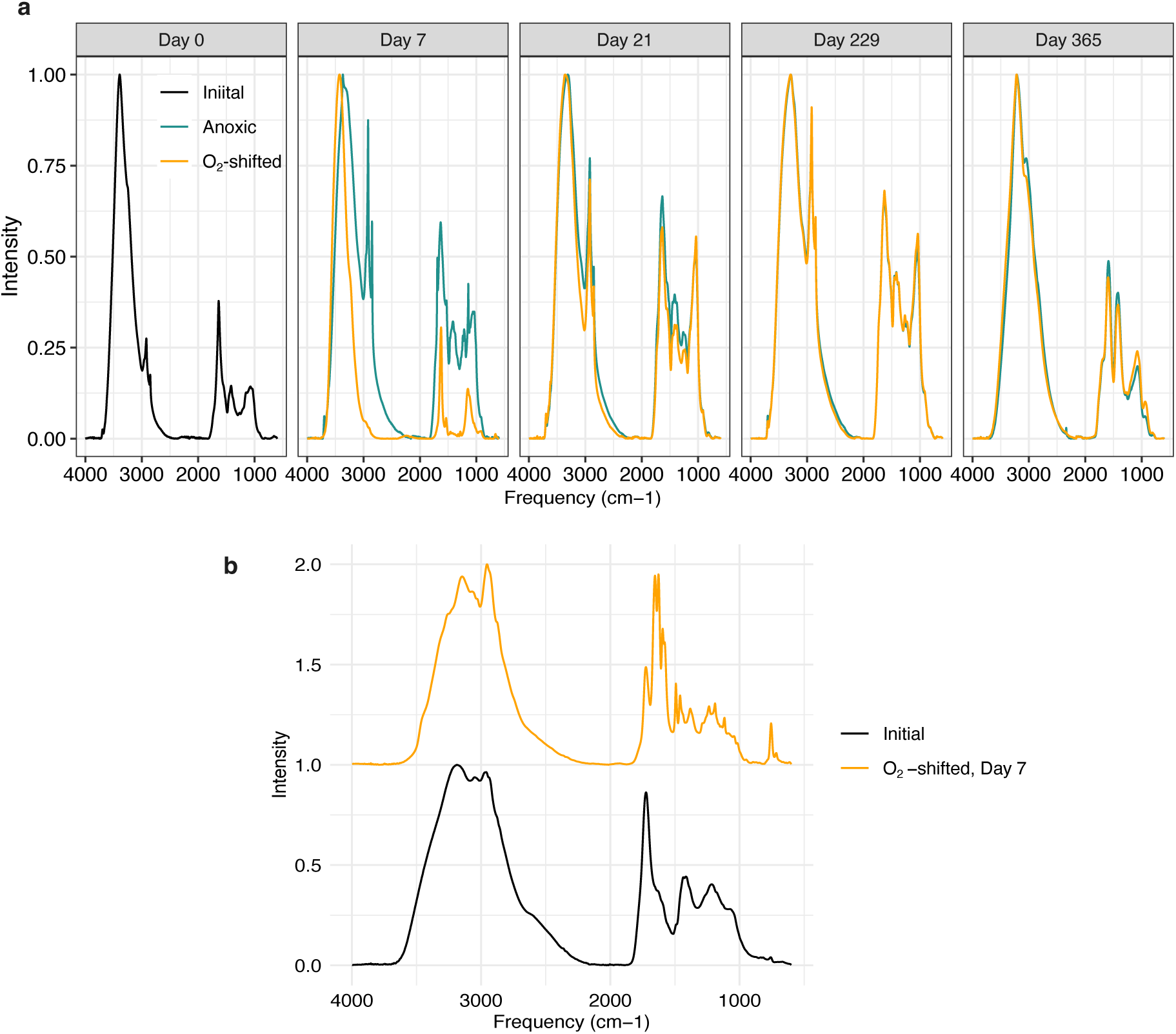
FTIR spectra of PB peat slurry extracts (**a**) and after desalting for the Day 7 samples The Day 7 samples were desalted and compared due to significant salt interferences (indicated by the strong O–H bending band at 1620 cm^-1^) in the O_2_-shifted sample (**a**).

**Figure S3:**
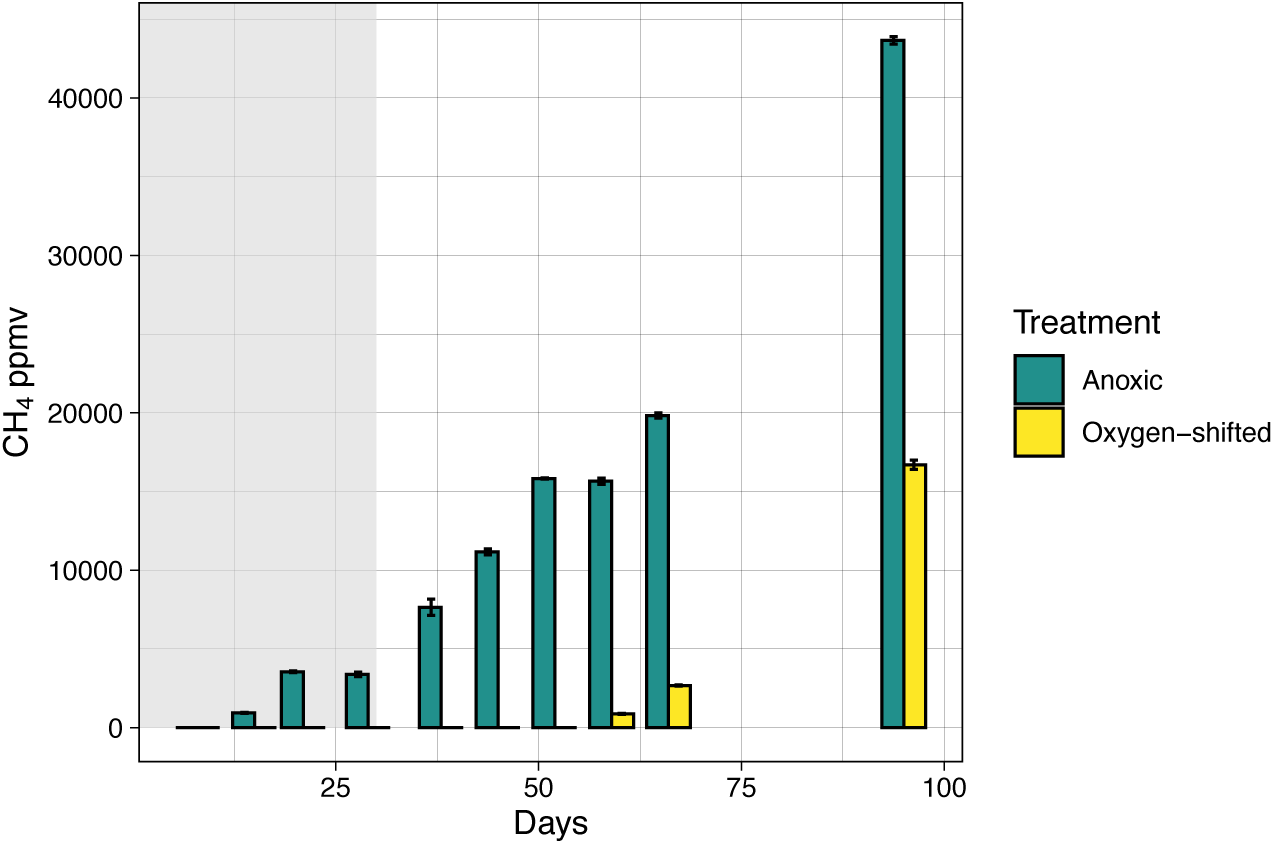
Headspace methane concentrations in the PB peat incubations when PB peat was exposed to a longer period of oxygenation (i.e., 4 weeks). Error bars are standard deviations around the mean of three replicates.

**Figure S4:**
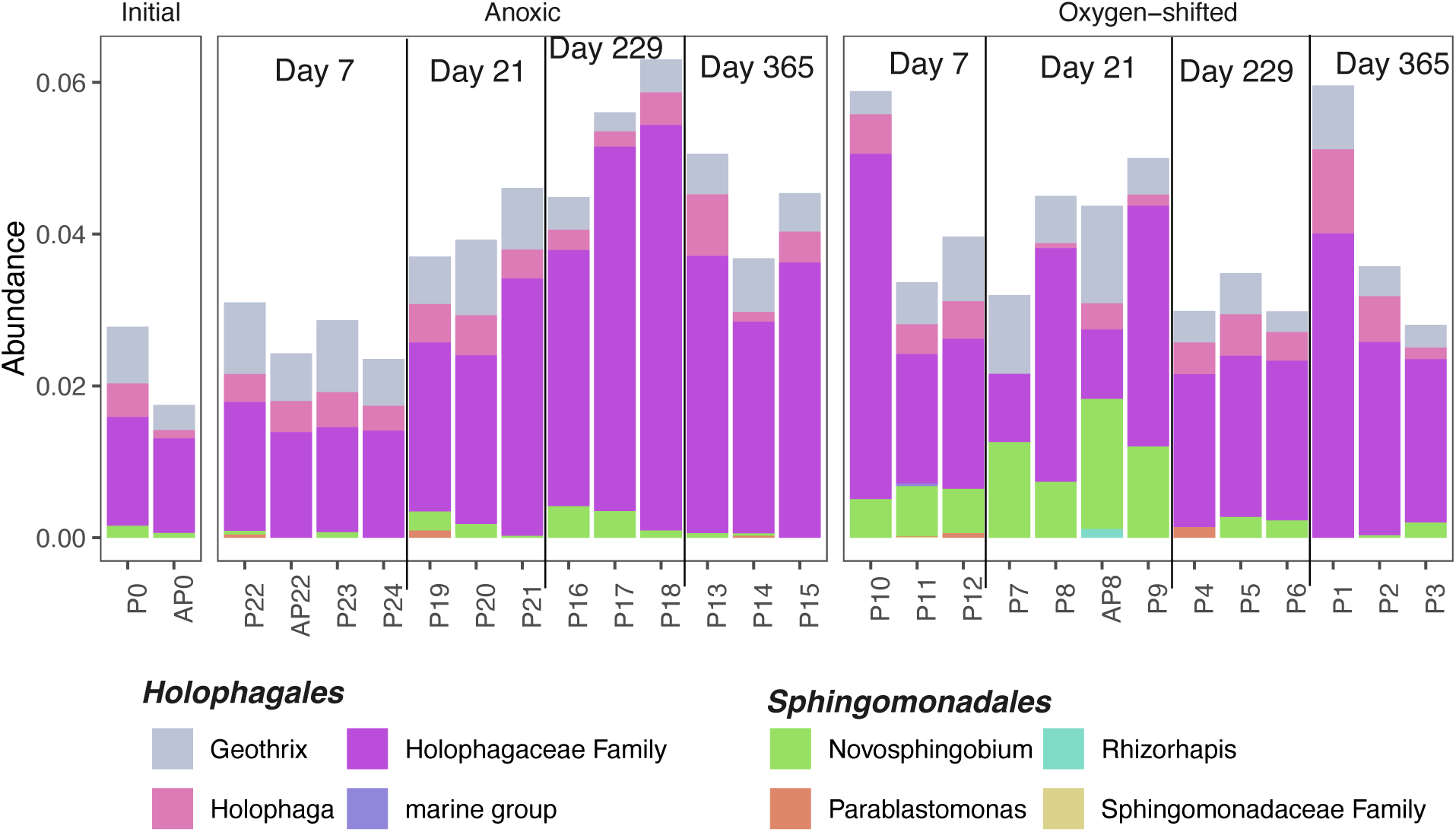
Relative abundance of selected lineages in the PB peat across timepoints. These lineages were identified as key indicator taxa for O_2_-stimulation of methanogenesis in the Ward Sphagnum peat.

**Figure S5:**
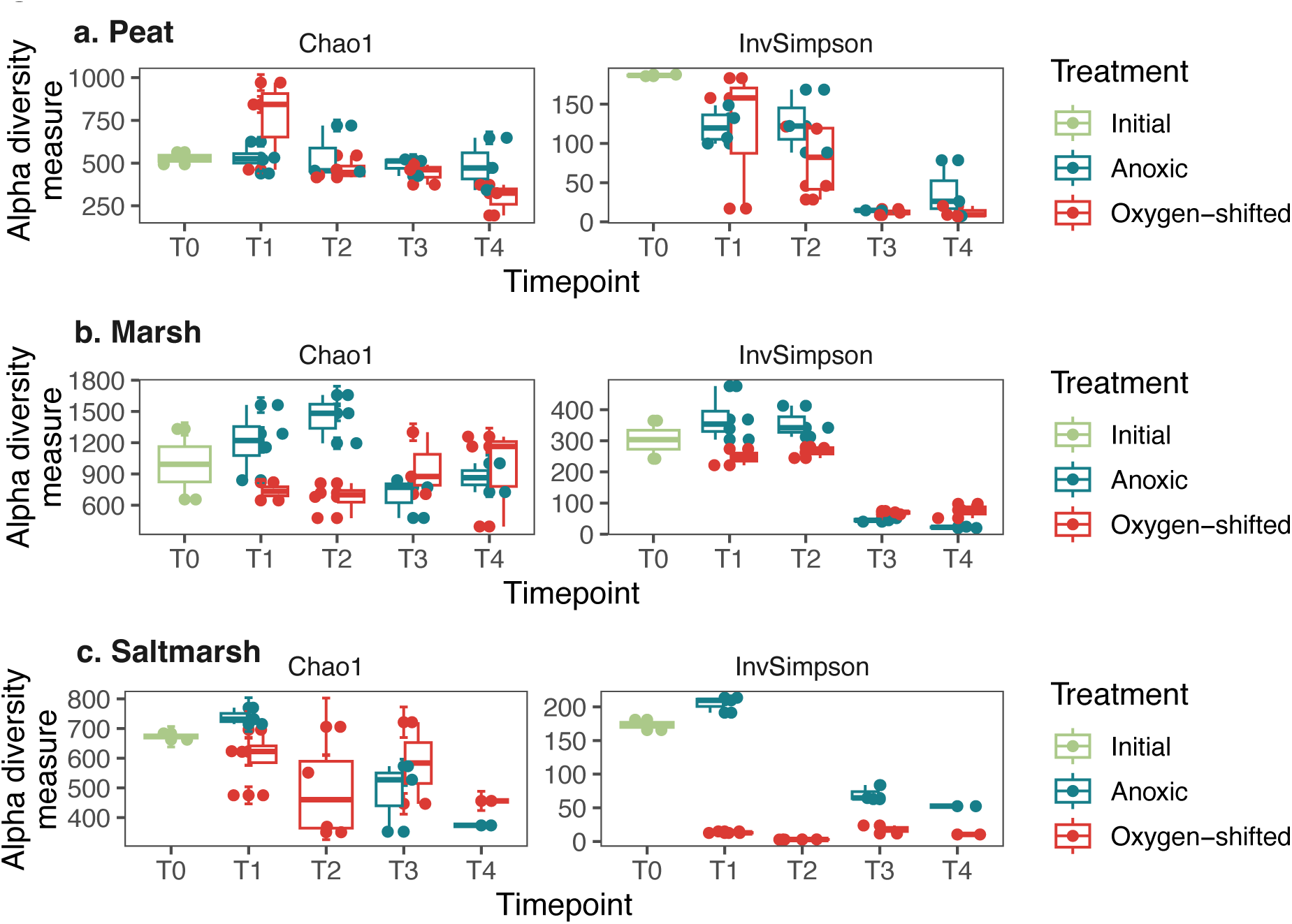
Estimates of microbial community alpha diversity (i.e., Observed number of ASVs and Inverse Simpson index) in the incubations across time for (a) peat, (b) marsh, and (c) saltmarsh. The filtered data used for alpha diversity estimation did not contain singletons. The data were rarefied to even depth before diversity estimation.

**Fig. S6:**
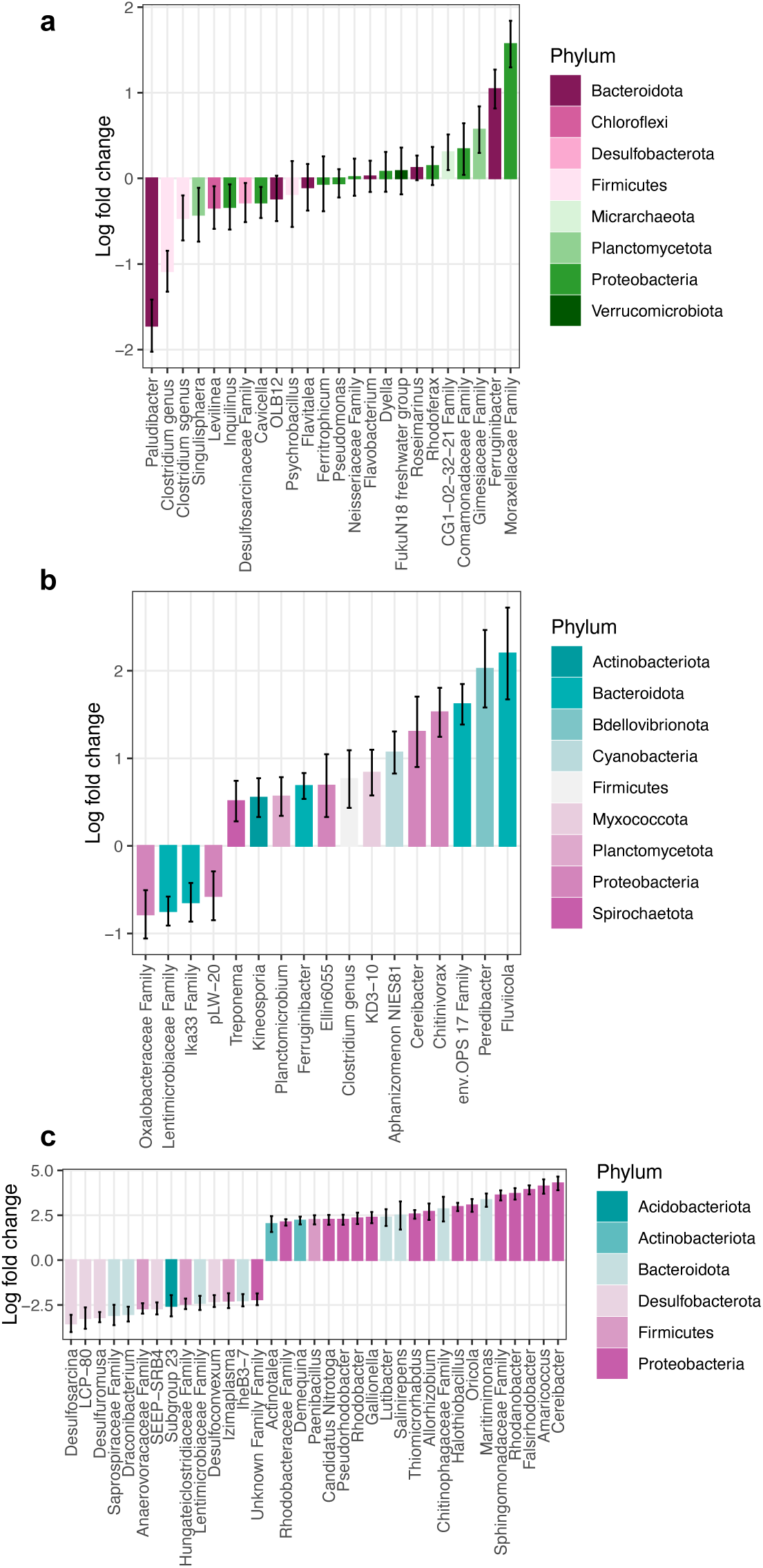
Summary of the differentially abundant ASVs between treatments in (**a**) PB peat, (**b**) FW marsh, and (**c**) saltmarsh. ANCOM-BC was used for differential abundance analysis. Since the microbiome changed substantially between treatments for saltmarsh, only those with a fold change of at least 2 are included in (**c**).

**Figure S7:**
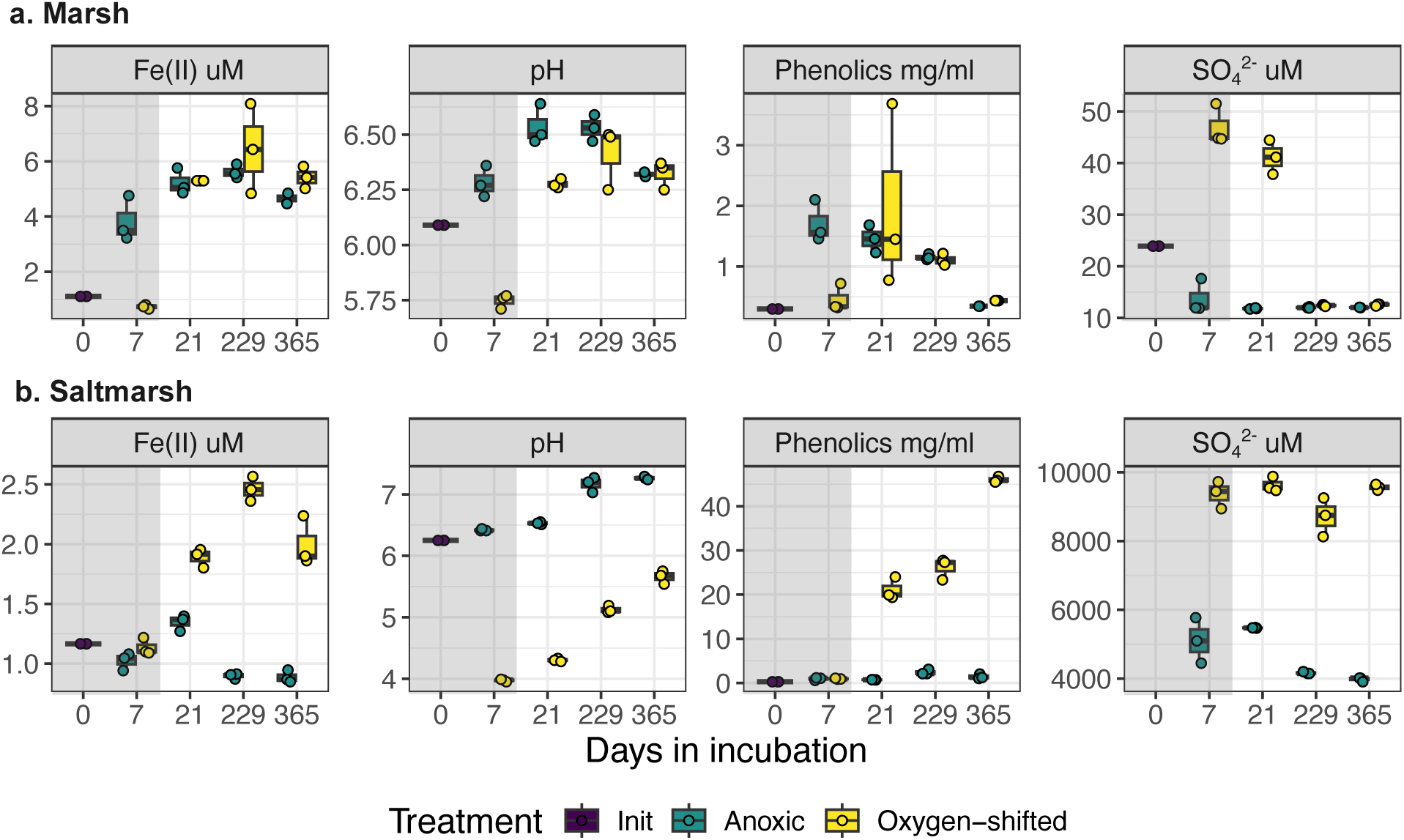
Key geochemical variables measured in FW marsh and saltmarsh incubations across timepoints in the experiment. Oxic period is indicated by the grey rectangles. Three replicate measurements are included in each boxplot (overlayed dots depict the actual values). Phenolics concentrations were measured as mg/ml gallic acid-equivalent.

## Notes

### Competing Interest Statement

The authors have declared no competing interest.

### Summary of Updates

Updated the author list and linked ORCIDs. Minor revisions to the abstract, and figures 3 & 5. This revision matches the version that has just been submitted for peer review.

